# A hybrid RNA FISH immunofluorescence protocol on *Drosophila* polytene chromosomes

**DOI:** 10.1101/2023.06.12.544616

**Authors:** Hannah E. Gilbonio, Gwyn L. Puckett, Erica Nguyen, Leila E. Rieder

**Affiliations:** Department of Biology, Emory University, Atlanta, GA, USA; Piedmont Virginia Community College, Charlottesville, VA, USA; Perelman School of Medicine, University of Pennsylvania, Philadelphia, PA, USA

**Keywords:** RNA FISH, IF, *Drosophila*, polytene chromosomes, transcription factor, histone genes, Mxc

## Abstract

**Objectives:** Investigating protein-DNA interactions is imperative to understanding fundamental concepts such as cell growth, differentiation, and cell development in many systems. Sequencing techniques such as ChIP-seq can yield genome-wide DNA binding profiles of transcription factors; however this assay can be expensive, time-consuming, may not be informative for repetitive regions of the genome, and depend heavily upon antibody suitability. Combining DNA fluorescence in situ hybridization (FISH) with immunofluorescence (IF) is a quicker and inexpensive approach which has historically been used to investigate protein-DNA interactions in individual nuclei. However, these assays are sometimes incompatible due to the required denaturation step in DNA FISH that can alter protein epitopes, hindering primary antibody binding. Additionally, combining DNA FISH with IF may be challenging for less experienced trainees. Our goal was to develop an alternative technique to investigate protein-DNA interactions by combining RNA FISH with IF.

**Results:** We developed a hybrid RNA FISH and IF protocol for use on *Drosophila melanogaster* polytene chromosome spreads in order to visualize colocalization of proteins and DNA loci. We demonstrate that this assay is sensitive enough to determine if our protein of interest, Multi-sex combs (Mxc), localizes to single-copy target transgenes carrying histone genes. Overall, this study provides an alternative, accessible method for investigating protein-DNA interactions at the single gene level in *Drosophila melanogaster* polytene chromosomes.

## INTRODUCTION

The relationships between a DNA locus and the proteins that target that locus affect fundamental processes such as DNA replication, transcription, and repair (1). A common technique used to investigate protein-DNA localization is ChIP-seq, which captures positional information of proteins across the genome. However, ChIP-seq involves several caveats: it is expensive, it requires access to sequencing platforms (2), and it is difficult to perform by inexperienced users. As an alternative to ChIP-seq, investigators utilize microscopy to reveal protein localization (3), monitor biochemical interactions between proteins and DNA (4), and quantify binding mechanisms that lead to the formation of protein-DNA complexes (5). To confirm whether a protein of interest is targeting a specific genomic locus, investigators have historically combined DNA fluorescent *in situ* hybridization (DNA FISH) with protein immunofluorescence (IF) (6,7). However, the protocol for DNA FISH is often incompatible with protein immunohistochemistry; it involves a DNA denaturation step that can reverse chemical crosslinks and denature protein epitopes, thus hindering primary antibody binding (8–10).

Combining DNA FISH with IF on *Drosophila* polytene chromosomes is a historically invaluable method for cytological analysis. Polytene chromosomes are formed through repetitive cycles of DNA endoreduplication without nuclear division. These polyploid cells can contain up to 1024 copies of the genome (11). This amplification of the genome gives rise to defined chromosomal banding patterns that represent chromatin regions and allows investigators to analyze relatively high-resolution protein binding patterns (12,13). Although researchers have applied DNA FISH and IF on polytenes in the past (6), the aforementioned incompatibility presents a need for an alternative to investigate protein-DNA interactions. To address this need, we developed a hybrid RNA FISH - IF method that indirectly marks genes on *Drosophila* polytene chromosomes.

Unlike DNA FISH, RNA FISH identifies the RNA transcripts that surround its parent gene (14), thus providing indirect genomic locus information. As RNA is single-stranded, RNA FISH does not require a denaturation step (15), which renders it more compatible with immunohistochemistry and more accessible for less-experienced trainees than DNA FISH.

We tested the ability of RNA FISH probes to mark genomic loci on polytene chromosomes, determined if single molecule RNA (smRNA) probes hybridize to local RNA or DNA, and optimized our hybrid RNA FISH and IF protocol. We used the *D. melanogaster* histone gene array as our model system. In the wildtype *D. melanogaster* genome, the endogenous histone locus consists of five histone genes arranged in a single histone array (HA). Arrays are tandemly repeated ∼107 times at a single locus (16). We also used engineered fly lines that carry transgenes with varying numbers of HAs, where even a single HA can attract specific transcription factors (17,18). Notably, this gene array titration allowed us to incrementally optimize the sensitivity of our RNA FISH - IF hybrid assay towards single-gene detection. We verified expected protein-DNA interactions on polytenes marked with histone smRNA FISH probes by immunostaining with Multi-sex combs (Mxc), a protein that only targets HAs (including both endogenous and transgenic HAs) (19–21). We sought to expand this technique for those working with any *Drosophila* transgene marked by reporters, specifically by leveraging smRNA probes targeting common markers like *mini-white*. However, we found that most single-copy genes did not give strong signal by smRNA FISH, suggesting that transcriptional level in salivary gland tissue contributes to detection limits. Overall, we present a protocol for investigating and visualizing protein binding at specific genomic locations that is inexpensive, quick, and accessible to *Drosophila* investigators at all levels.

## METHODS

### Fly stocks

We maintained all *Drosophila melanogaster* fly lines on standard cornmeal-molasses food at 18°C. We used third instar larvae for dissections. We obtained fly stocks from Bloomington *Drosophila* Stock Center (*y,w*: stock #1495) or stocks as gifts from the Duronio and Marzluff laboratories (17,22).

### Antibodies and RNA FISH probes

We used the following antibodies: guinea pig anti-Mxc (1:5000; gift from Duronio and Marzluff laboratories); Goat anti-Guinea Pig AlexaFluor 488 (1:1000) (Invitrogen #A-11073); Goat anti-Guinea Pig AlexaFluor 647 (1:1000) (Invitrogen #A-21450). We obtained custom RNA FISH probe sets from LGC Biosearch Technologies using the Stellaris Probe Designer version 4.2. All RNA FISH probes used in this procedure were coupled to Quasar 570 or Quasar 670 fluorophores.

### RNase decontamination

We made RNase-free reagents from DEPC-treated water and RNase-free 1X PBS (23). We used RNase AWAY (Spectrum Chemical MFG Corp #970-66898) to clean all surfaces and tools and used filtered pipette tips. We cleaned slide holders by washing them in a mixture of DEPC-treated water and RNase AWAY.

### Sample preparation

We dissected salivary glands from third instar larvae and fixed the glands containing polytene chromosomes with three separate fixatives (Fix 1: 4% paraformaldehyde, 1% Triton X-100 in RNase-free 1X PBS) (Fix 2: 4% paraformaldehyde, 50% acetic acid in DEPC-treated water) (Fix 3: 1:2:3 lactic acid:DEPC-treated water:glacial acetic acid) (18) using DEPC-treated reagents and RNase-free materials. We transferred salivary glands to the 1:2:3 fixative (∼20 uL) on a siliconized (RainEX) 22mm square coverslip surface. We placed a glass slide on the coverslip containing the salivary glands and third fix and quickly flipped over. We froze slides in liquid nitrogen and immediately removed their coverslips. We briefly stored slides in RNase-free 1X PBS and immediately proceeded with immunohistochemistry.

### Immunohistochemistry

We washed slides in 1% Triton X-100 in RNase-free 1X PBS and rocked gently for 10 minutes. We washed slides twice for 5 minutes in RNase-free 1X PBS and marked the sample perimeter of each slide with an ImmEdge pen (Vector Laboratories #H-4000) in between both washes. We added 250 uL of blocking solution (0.5% UltraPure BSA (Invitrogen #AM2616) in RNase-free 1X PBS) to the sample area of each slide and incubated them at room temperature in a dark humid chamber for 1 hour, shaking gently. Using coverslips, we then incubated slides with 40 uL of diluted primary antibody in blocking solution and incubated them overnight in a dark, sealed, humid chamber at 4°C. We washed slides 3 times for 5 minutes in RNase-free 1X PBS. We applied 40ul of diluted secondary antibody in blocking solution to each slide (including coverslips) and incubated them for 2 hours in a dark humid chamber at room temperature. We washed slides 3 times for 5 minutes in RNase-free 1X PBS. We added 250 uL of post-fixative (4% paraformaldehyde in RNase-free 1X PBS) to each sample area for 3 minutes at room temperature. We then washed each slide 3 times for 5 minutes in RNase-free 1X PBS.

### RNA FISH

We washed slides for 5 minutes in ∼30 mL Wash Buffer (1:10:1 20x SSC:DEPC-treated water:Deionized Formamide) at room temperature. We added 125 uL of Hybridization Buffer (100 mg/mL dextran sulfate and 10% formamide in 2x SSC and DEPC-treated water) and diluted RNA FISH probe (5 uL of 25uM probe:120 uL Hybridization Buffer) to slides (including coverslips) which were incubated in a dark, humid, sealed chamber at 37°C overnight (∼16 hours). We washed slides twice for 10 minutes in prewarmed (37°C) Wash Buffer in the dark. We then added 250 uL Wash Buffer and diluted DAPI (25 ng/mL DAPI) onto the sample area and incubated slides in a dark, humid chamber at 37°C for 30 minutes. We mounted slides with

∼15uL of VECTASHIELD Antifade Mounting Medium (VWR #101098-042) and coverslips. We sealed the samples with nail polish. We immediately proceeded to imaging.

### Microscopy

We used a widefield Zeiss AXIO Scope A1 microscope with X-Cite 120 LED Boost fluorescent light source and a 40x Plan-neofluar 0.75 NA objective paired with ZEN 3.6 (blue edition). We visualized .czi files with ImageJ Version 1.53t.

### RNase treatment

After dissecting salivary glands, we treated the glands with 0.1% Triton X-100 for 2 minutes prior to adding 100 ug/mL RNase A (NEB #T3018L) and performed a 1 hour incubation at RT (24) before beginning fixation.

## RESULTS

A high concentration of transcripts surrounds the parent gene locus in many cells. We therefore hypothesized that RNA FISH would mark genetic loci on *Drosophila* polytene chromosomes.

We performed RNA FISH on wild-type polytene chromosomes using smRNA FISH probe sets against either *histone2b* or *histone3* transcripts because they are the longest histone genes. The endogenous histone locus, which carries ∼100 tandem histone gene arrays (16), is located on chromosome 2L near the centromere. Our *histone2b* and *histone3* RNA FISH probe sets effectively targeted this region (**Fig. 1A**). Since most genes do not exist in multiple copies, we next performed the same RNA FISH assay on transgenic lines carrying HA transgenes, either with 12 tandem copies of the histone gene array or a single array. Our *histone2b* and *histone3* probe sets effectively detected these transgenes (**Fig. 1B-C**).

**Figure 1.**
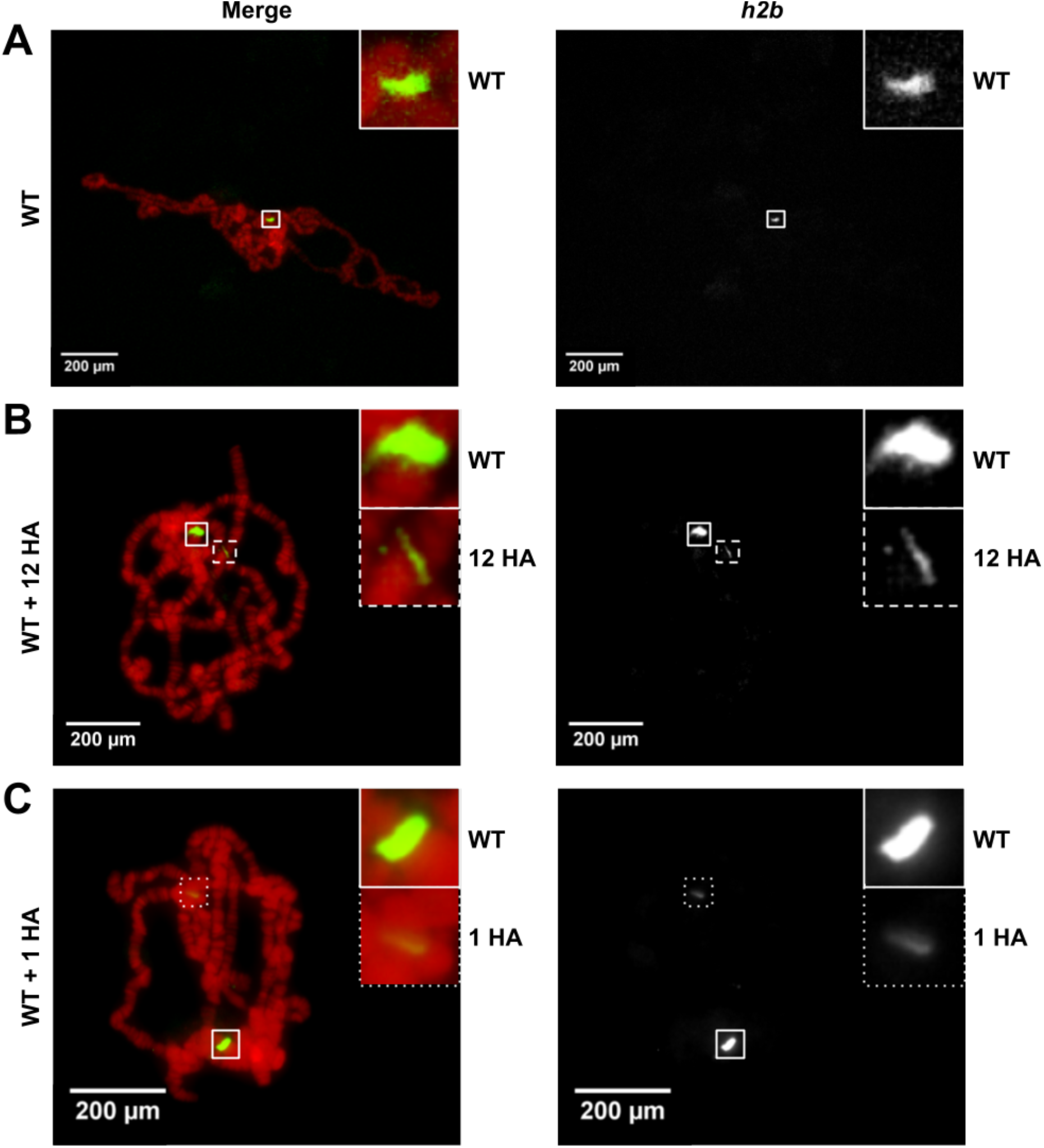
RNA FISH (*h2b*) on **A** wildtype, **B** wildtype with a 12 copy histone array transgene, and **C** wildtype with a single copy histone array transgene on *D. melanogaster* polytene chromosomes.

To confirm that our RNA FISH probe sets are detecting histone loci, and to confirm that our RNA FISH protocol is compatible with immunofluorescence (IF), we combined our RNA FISH assay with antibody detection of the histone-locus specific factor Multi-sex combs (Mxc) (22,25). Mxc also localizes to transgenes carrying histone gene arrays on polytene chromosomes (17,18,22). We observed that Mxc signal colocalizes with *histone2b* and *histone3* RNA FISH, confirming these locations as the endogenous histone locus (**Fig. 2A**) and the transgenic loci (**Fig. 2B-C**). While developing this protocol, we intentionally conducted IF first because certain RNA FISH regents can alter protein epitopes. We included a postfixative step in between IF and RNA FISH to preserve the IF signal (23), included larger-volume washes, and increased the concentration of probe to 3 uL of 25 uM probe per 100 uL. All of these steps contributed to boosting RNA FISH signals when combined with IF.

**Figure 2.**
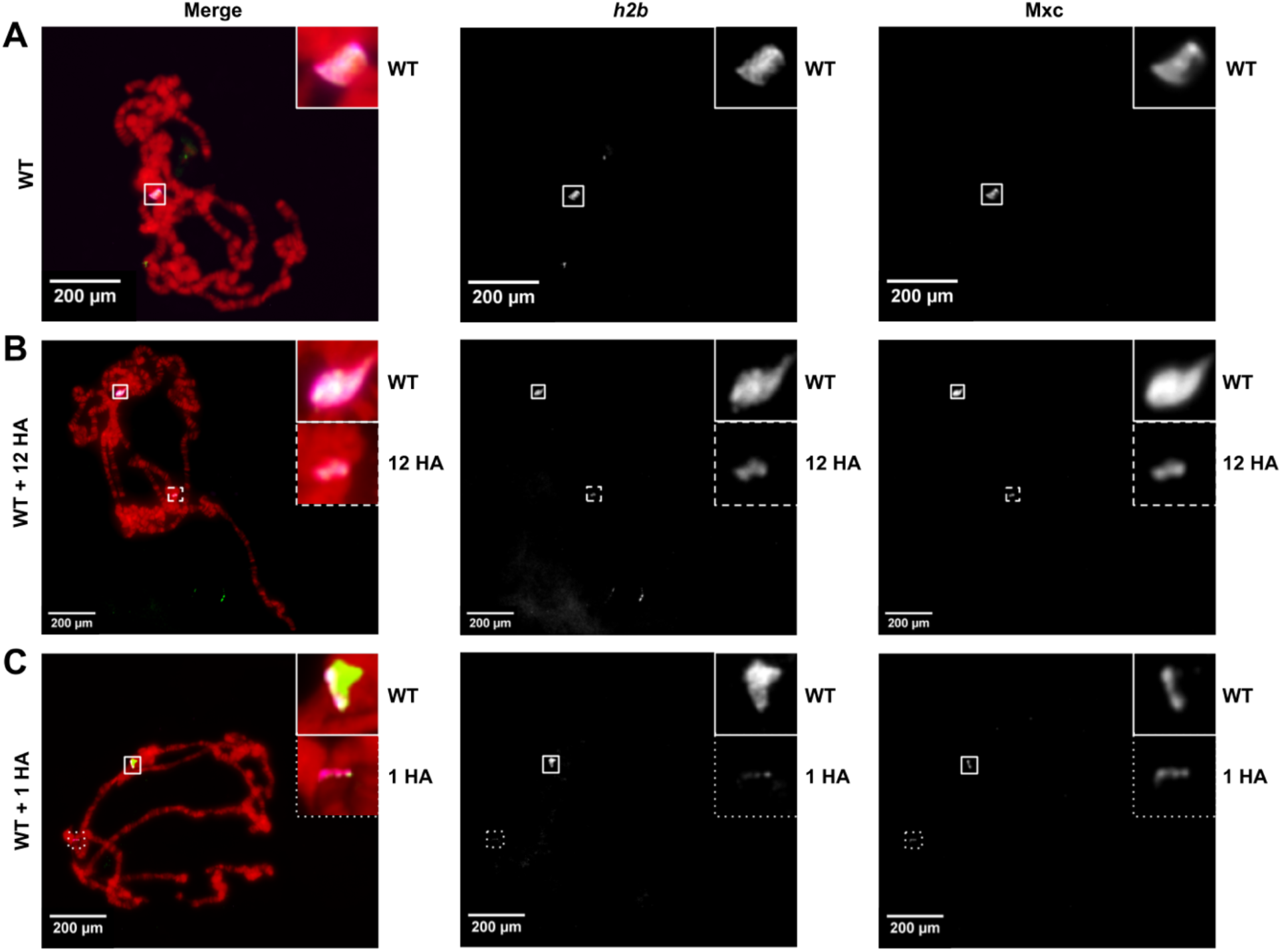
RNA FISH (*h2b*) and IF (Mxc) on **A** wildtype, **B** wildtype with a 12 copy histone array transgene, and **C** wildtype with a single copy histone array transgene on *D. melanogaster* polytene chromosomes.

Given that we verified Mxc colocalization with histone genes, we wanted to further test the ability of both assays to mark different single-copy genomic loci on polytenes. We tested our procedure on *mini-white*, a common transgene marker that we use to mark HA transgenes on chromosome 3R, as well as *roX2*, an X-linked long non-coding RNA that is only expressed in males. *RoX2* participates in *Drosophila* dosage compensation and coats the male X-chromosome (26,27). We did not detect a signal for the *mini-white* gene **(Fig. 3A)**. While we clearly detected *roX2* on male X-chromosomes **(Fig. 3B)**, as previously documented, we saw no RNA FISH signals for the *roX2* locus in female polytenes **(Fig. 3C)**. To verify that our RNA FISH probes sets were binding to mRNA transcripts and not binding directly to DNA, we introduced RNase A into the procedure. We observed a large reduction in signal **(Fig. 3D-E)**, suggesting that our RNA FISH probes are binding directly to local mRNA.

**Figure 3.**
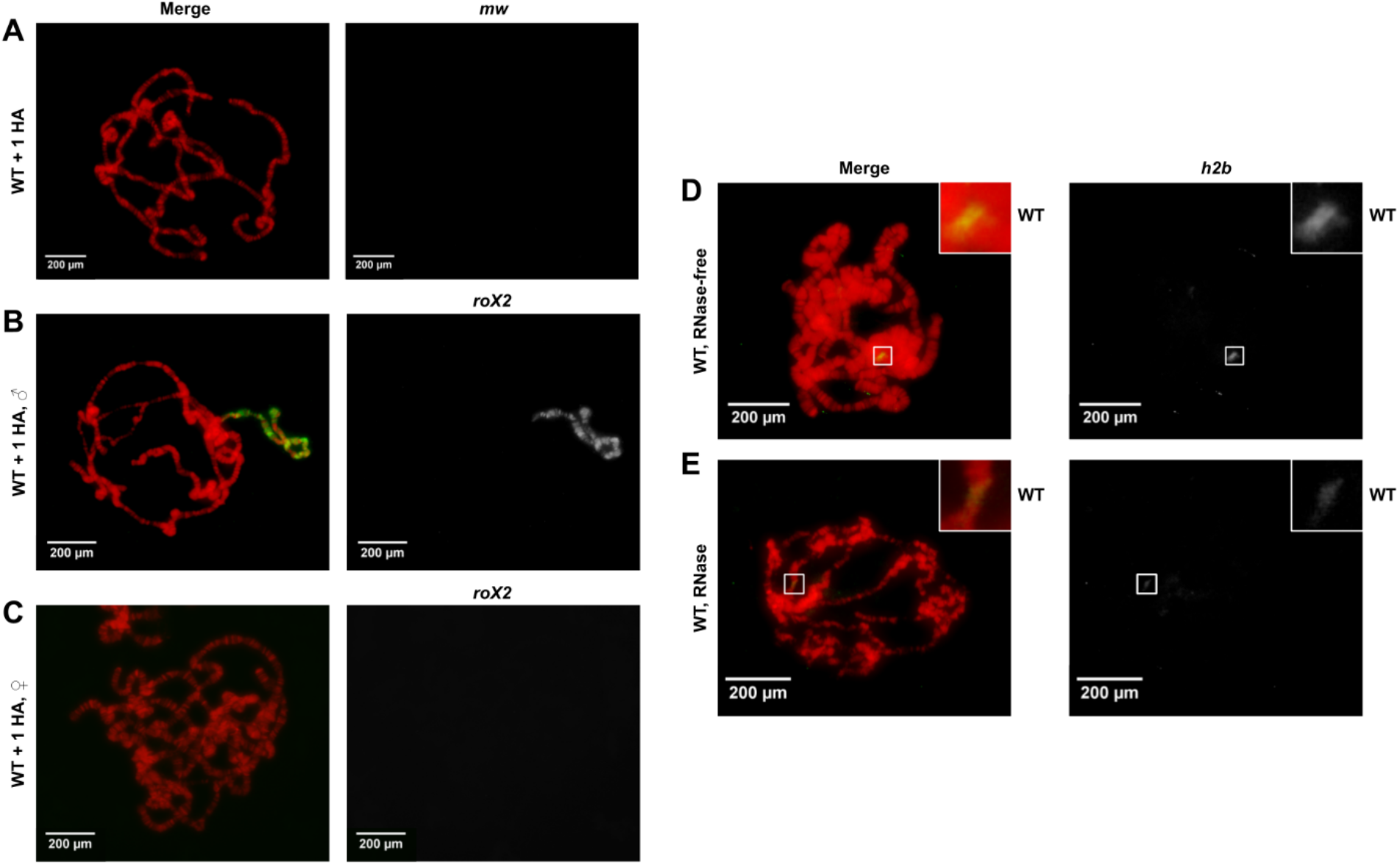
RNA FISH on wildtype *D. melanogaster* polytene chromosomes with a single copy histone array transgene using **A** *mw*. **B** *roX2* on males, and **C** *roX2* on females. RNA FISH (*h2b*) on wildtype polytenes **D** without RNase treatment and **E** with RNase treatment.

## DISCUSSION

Our goal in developing a hybrid RNA FISH and IF protocol for polytene chromosomes was to create an alternative method to visualize protein-DNA interactions in *Drosophila*. Here, we successfully combined RNA FISH and IF to visualize Mxc (a protein that only targets histone genes) colocalizing with histone gene loci. This proof of principle suggests that our hybrid assay could be applied to other proteins and genomic loci of interest. Unfortunately, we found that *mini-white*, a common transgene marker in *Drosophila*, was not visible by RNA FISH on polytene chromosomes. Considering that the wild-type *white* gene has very low expression levels in salivary glands (28), we are not surprised by the absence of *mini-white* signal. These observations suggest that our RNA FISH technique is more applicable for visualizing loci of genes that are highly expressed in larval salivary glands. Due to the relatively low cost of the protocol and reagents in addition to the 3-day turnaround time, we believe this hybrid RNA FISH and IF procedure is an accessible method for testing protein-DNA colocalization, especially for those with limited wet lab experience.

## LIMITATIONS

This protocol is most likely limited to investigating genes in *Drosophila* polytene chromosomes that have high expression in *Drosophila* larval salivary glands. We only tested this method in *Drosophila melanogaster*, yet we see no reason why this assay cannot be applied to other *Drosophila* species. Polytene chromosomes represent up to 1024 copies of the genome (11); we did not test our protocol on diploid cells. However, as others have observed that RNA FISH signals often cluster around genomic loci, RNA FISH + IF is likely a viable technique to simultaneously visualize DNA loci and proteins, even in diploid cells.

## ABBREVIATIONS

smRNA: single molecule RNA
FISH: fluorescent in situ hybridization
IF: immunofluorescence
Mxc: Multi sex-combs
ChIP-seq: chromatin immunoprecipitation sequencing
HA: histone array
DEPC: diethyl pyrocarbonate

## DECLARATIONS

## Acknowledgements

We thank the Lerit and Deal labs at Emory University and the Larschan lab at Brown University for sharing reagents, the Duronio and Marzluff labs at UNC Chapel Hill for antibodies and fly stocks, and other members of the Rieder lab for experiment and manuscript feedback.

## Author’s contributions

LER conceived the project. EN and GP obtained preliminary data. HEG conducted the experiments and wrote the manuscript. All authors edited the manuscript. LER supervised the project.

## Funding

Financial support was provided by R35GM142724 to LER and R35GM142724-01S1 supplement to HEG.

Data availability

Data are available from the corresponding author upon request.

## Ethics approval and consent to participate

Not applicable.

## Consent for publication

Not applicable.

## References

1. Cozzolino F, Iacobucci I, Monaco V, Monti M. Protein-DNA/RNA Interactions: An Overview of Investigation Methods in the -Omics Era. J Proteome Res. 2021 Jun 4;20(6):3018–30.

2. Park PJ. ChIP-Seq: advantages and challenges of a maturing technology. Nat Rev Genet. 2009 Oct;10(10):669.

3. Beckwitt EC, Kong M, Van Houten B. Studying protein-DNA interactions using atomic force microscopy. Semin Cell Dev Biol. 2018 Jan;73:220–30.

4. Crickard JB. Single Molecule Imaging of DNA-Protein Interactions Using DNA Curtains. Methods Mol Biol. 2023;2599:127–39.

5. Ritzefeld M, Walhorn V, Anselmetti D, Sewald N. Analysis of DNA interactions using single-molecule force spectroscopy. Amino Acids. 2013 Jun;44(6):1457–75.

6. Rudkin GT, Stollar BD. High resolution detection of DNA-RNA hybrids in situ by indirect immunofluorescence. Nature. 1977 Feb 3;265(5593):472–3.

7. Yu Z, Potapova TA. Superresolution Microscopy for Visualization of Physical Contacts Between Chromosomes at Nanoscale Resolution. Methods Mol Biol. 2022;2458:359–75.

8. Cremer M, Grasser F, Lanctôt C, Müller S, Neusser M, Zinner R, et al. Multicolor 3D Fluorescence In Situ Hybridization for Imaging Interphase Chromosomes. In: Hancock R, editor. The Nucleus: Volume 1: Nuclei and Subnuclear Components. Totowa, NJ: Humana Press; 2008. p. 205–39.

9. Fowler CB, Evers DL, O’Leary TJ, Mason JT. Antigen retrieval causes protein unfolding: evidence for a linear epitope model of recovered immunoreactivity. J Histochem Cytochem. 2011 Apr;59(4):366–81.

10. Chaumeil J, Micsinai M, Skok JA. Combined immunofluorescence and DNA FISH on 3D-preserved interphase nuclei to study changes in 3D nuclear organization. J Vis Exp. 2013 Feb 3;(72):e50087.

11. Lindsley DL, Zimm GG. The Genome of Drosophila Melanogaster. Academic Press; 2012. 1133 p.

12. Ashburner M, Golic KG, Scott Hawley R. Drosophila: A Laboratory Handbook. Cold Spring Harbor Laboratory Press; 2005. 1409 p.

13. Lavrov S, Déjardin J, Cavalli G. Combined Immunostaining and FISH Analysis of Polytene Chromosomes. In: Henderson DS, editor. Drosophila Cytogenetics Protocols. Totowa, NJ: Humana Press; 2004. p. 289–303.

14. Young AP, Jackson DJ, Wyeth RC. A technical review and guide to RNA fluorescence in situ hybridization. PeerJ. 2020 Mar 19;8:e8806.

15. Greenberg E, Hochberg-Laufer H, Blanga S, Kinor N, Shav-Tal Y. Cytoplasmic DNA can be detected by RNA fluorescence in situ hybridization. Nucleic Acids Res. 2019 Oct 10;47(18):e109.

16. Bongartz P, Schloissnig S. Deep repeat resolution-the assembly of the Drosophila Histone Complex. Nucleic Acids Res. 2019 Feb 20;47(3):e18.

17. Salzler HR, Tatomer DC, Malek PY, McDaniel SL, Orlando AN, Marzluff WF, et al. A sequence in the Drosophila H3-H4 Promoter triggers histone locus body assembly and biosynthesis of replication-coupled histone mRNAs. Dev Cell. 2013 Mar 25;24(6):623–34.

18. Rieder LE, Koreski KP, Boltz KA, Kuzu G, Urban JA, Bowman SK, et al. Histone locus regulation by the Drosophila dosage compensation adaptor protein CLAMP. Genes Dev. 2017 Jul 15;31(14):1494–508.

19. Ma T, Van Tine BA, Wei Y, Garrett MD, Nelson D, Adams PD, et al. Cell cycle-regulated phosphorylation of p220(NPAT) by cyclin E/Cdk2 in Cajal bodies promotes histone gene transcription. Genes Dev. 2000 Sep 15;14(18):2298–313.

20. White AE, Burch BD, Yang XC, Gasdaska PY, Dominski Z, Marzluff WF, et al. Drosophila histone locus bodies form by hierarchical recruitment of components. J Cell Biol. 2011 May 16;193(4):677–94.

21. Zhao J, Kennedy BK, Lawrence BD, Barbie DA, Matera AG, Fletcher JA, et al. NPAT links cyclin E–Cdk2 to the regulation of replication-dependent histone gene transcription. Genes Dev. 2000 Sep 15;14(18):2283–97.

22. Koreski KP, Rieder LE, McLain LM, Chaubal A, Marzluff WF, Duronio RJ. Drosophila histone locus body assembly and function involves multiple interactions. Mol Biol Cell. 2020 Jul 1;31(14):1525–37.

23. Kochan J, Wawro M, Kasza A. Simultaneous detection of mRNA and protein in single cells using immunofluorescence-combined single-molecule RNA FISH. Biotechniques. 2015 Oct;59(4):209–12, 214, 216 passim.

24. Singh AK, Choudhury SR, D. S, Zhang J, Kissane S, Dwivedi V, et al. The RNA helicase UPF1 associates with mRNAs co-transcriptionally and is required for the release of mRNAs from gene loci. Elife [Internet]. 2019 Mar 25;8. Available from: http://dx.doi.org/10.7554/eLife.41444

25. Terzo EA, Lyons SM, Poulton JS, Temple BRS, Marzluff WF, Duronio RJ. Distinct self-interaction domains promote Multi Sex Combs accumulation in and formation of the Drosophila histone locus body. Mol Biol Cell. 2015 Apr 15;26(8):1559–74.

26. Kelley RL, Meller VH, Gordadze PR, Roman G, Davis RL, Kuroda MI. Epigenetic spreading of the Drosophila dosage compensation complex from roX RNA genes into flanking chromatin. Cell. 1999 Aug 20;98(4):513–22.

27. Meller VH, Gordadze PR, Park Y, Chu X, Stuckenholz C, Kelley RL, et al. Ordered assembly of roX RNAs into MSL complexes on the dosage-compensated X chromosome in Drosophila. Curr Biol. 2000 Feb 10;10(3):136–43.

28. Pirrotta V, Bickel S, Mariani C. Developmental expression of the Drosophila zeste gene and localization of zeste protein on polytene chromosomes. Genes Dev. 1988 Dec;2(12B):1839–50.

